# Lateralised cerebral processing of abstract linguistic structure in clear and degraded speech

**DOI:** 10.1101/2020.02.05.934604

**Authors:** Qingqing Meng, Yiwen Li Hegner, Iain Giblin, Catherine McMahon, Blake W Johnson

## Abstract

Providing a plausible neural substrate of speech processing and language comprehension, cortical activity has been shown to track different levels of linguistic structure in connected speech (syllables, phrases and sentences), independent of the physical regularities of the acoustic stimulus. In the current study, we investigated the effect of speech intelligibility on this brain activity as well as the underlying neural sources. Using magnetoencephalography (MEG), brain responses to natural speech and noise-vocoded (spectrally-degraded) speech in nineteen normal hearing participants were measured. Results showed that cortical MEG coherence to linguistic structure changed parametrically with the intelligibility of the speech signal. Cortical responses coherent with phrase and sentence structures were left-hemisphere lateralized, whereas responses coherent to syllable/word structure were bilateral. The enhancement of coherence to intelligible compared to unintelligible speech was also left lateralized and localized to the parasylvian cortex. These results demonstrate that cortical responses to higher level linguistics structures (phrase and sentence level) are sensitive to speech intelligibility. Since the noise-vocoded sentences simulate the auditory input provided by a cochlear implant, such objective neurophysiological measures have potential clinical utility for assessment of cochlear implant performance.

## Introduction

Neural oscillations in the delta, theta and gamma frequency bands have been hypothesized as important mechanisms for speech perception. Recent neurolinguistic models (Giraud & Poeppel, 2012; Hickok & Poeppel, 2007) have proposed that these brain rhythms serve to segregate and package linguistic units at different time scales (corresponding to the prosodic, syllabic and phonemic time scales in speech) for further processing. A number of studies have reported that the auditory cortex exhibits activity that becomes phase-synchronized to the speech temporal envelope (Ahissar et al., 2001; Lakatos et al., 2005; Luo & Poeppel, 2007; Peelle, Gross, & Davis, 2013; Zion Golumbic et al., 2013). Prominent quasi-periodic cues occur at the syllabic rate of speech (corresponding to the speech envelope), which has a rate of about 4-7 Hz in natural speech (Chandrasekaran, Trubanova, Stillittano, Caplier, & Ghazanfar, 2009; MacNeilage, 1998). Psychophysical studies (Drullman, Festen, & Plomp, 1994; Shannon, Zeng, Kamath, Wygonski, & Ekelid, 1995) have demonstrated that the speech envelope is a powerful cue for speech perception and is also important for comprehension of speech processed by cochlear implants (Shannon, Fu, Galvin, & Friesen, 2004). Often described as “cortical entrainment” (Ding & Simon, 2014), neural phase-locking to syllable rate modulations of the speech envelope has been suggested to serve as a mechanism for the perceptual segmentation of the continuous speech stream into meaningful chunks, a parsing that facilitates extraction of linguistic information (Luo & Poeppel, 2007).

An important line of evidence for this proposition comes from studies showing that cortical phase-locking is significantly attenuated when speech intelligibility is degraded by destroying fine-structure while preserving the overall speech envelope (Peelle et al., 2013; Ding, Chatterjee, & Simon, 2014; Rimmele, Zion Golumbic, Schröger, & Poeppel, 2015). This line of reasoning is contentious, however. Other studies have argued that the speech entrainment phenomenon is largely or entirely determined by the acoustic, rather than linguistic, features of speech signal (Howard & Poeppel, 2010; Doelling, Arnal, Ghitza, & Poeppel, 2014; Millman, Johnson, & Prendergast, 2015). As discussed in several reviews (Peelle & Davis, 2012; Ding & Simon, 2014; Zoefel & VanRullen, 2015), this debate has arisen largely because it is difficult to unambiguously disentangle speech intelligibility and speech acoustics in these experiments.

A recent magnetoencephalography (MEG) study provides an important methodological advance by demonstrating that auditory cortical activity can track *abstract* linguistic structures, i.e., linguistic regularities that are embedded in connected speech but have no physical presence in the acoustic properties of the signal (Ding, Melloni, Zhang, Tian, & Poeppel, 2016). By presenting short sentences constructed with the same syntactic structure to participants in an isochronous manner, concurrent cortical tracking activity to syllable, phrase and sentence level linguistic structures was reported. Importantly, this neural tracking activity of the larger linguistic structures at phrase and sentence level is unambiguously dissociated from any acoustic cues to these units, as there are no physical phrase or sentence boundaries present in the isochronous speech signal. The authors concluded that a grammar-based internal construction process corresponding to the hierarchical linguistic structure must have been carried out (Ding, Melloni, Tian, & Poeppel, 2017).

In the current study we used the experimental paradigm of Ding et al. (2016) to investigate how speech intelligibility affects neural tracking of the speech stream, measured with MEG. Unlike previous studies, the Ding et al. (2016) paradigm permits an unambiguous separation of linguistic and acoustic cues; and further, permits the comparison of intelligibility effects on neural responses associated with distinct timescales (syllable, phrase and sentence) in connected speech. A parametric reduction in speech intelligibility was achieved using noise-vocoding (Shannon et al., 1995), which progressively reduces the amount of spectral detail present in the speech signal (i.e. the number of frequency channels used in the vocoding) while closely preserving the temporal envelope.

## Materials & Methods

### Participants

Experimental participants were 19 native speakers of English aged between 18 to 38 years old (mean 25 years old; 12 female) with normal hearing and no history of neurological, psychiatric, or developmental disorders (self-reported). All participants were right-handed and gave written informed consent under the process approved by the Human Subjects Ethics Committee of Macquarie University.

### Stimuli

All speech materials were synthesized using the MacinTalk text to speech synthesizer (male voice Alex, Mac OS X 10.11.4). In total, 180 four-syllable (a monosyllabic word for each syllable) English sentences were generated to form a sentence list (**Supplementary Material**). All sentences in the list followed the same syntactic structure: adjective/pronoun + noun + verb + noun. Each syllable was synthesized independently, and all the synthesized syllables (200 – 376 ms in duration) were adjusted to 320 ms by truncation or padding silence at the end. The offset of each syllable was smoothed with a 25-ms cosine window.

From the 180 sentences in the total pool, 60 (first set) were randomly selected to be presented in the unprocessed form (“natural speech”). The same set of 60 sentences were used to generate “shuffled sentences.” A second set of 60 sentences were randomly selected from the remaining 120 sentences for “16 channel noise vocoding”, and the remaining set (third set) of 60 sentences were used for “8 channel noise vocoding”.

Sentences were presented in a trial consisting of 12 sentences of the same type. From the sub pools of 60 sentences/condition, 12 sentences were randomly drawn from the pool of 60 for the first trial, another 12 randomly drawn from the remainder of 48, and so on to produce 5 trials of 12 sentences for each condition. This process was repeated six times to produce a total of 30 trials for each condition for the whole experiment. Over the 30 trials, each sentence was repeated six times. In each trial, 12 sentences were presented isochronously. (We note that this method of stimulus construction and presentation results in speech that is significantly less intelligible than more naturalistic (non-isochronous) speech; Smith, Delgutte, & Oxenham, 2002; Ding et al., 2014).

In catch trials (“outlier” trial in the terminology from Ding et al., 2016), 3 consecutive words selected from a random position within a trial, were replaced by three random words to abolish any meaningful sentence structure. There were eight outlier trials for each intelligibility condition.

A schematic plot of the linguistic units embedded in the isochronously presented syllable streams is depicted in **Figure 1** below:

**Figure 1:**
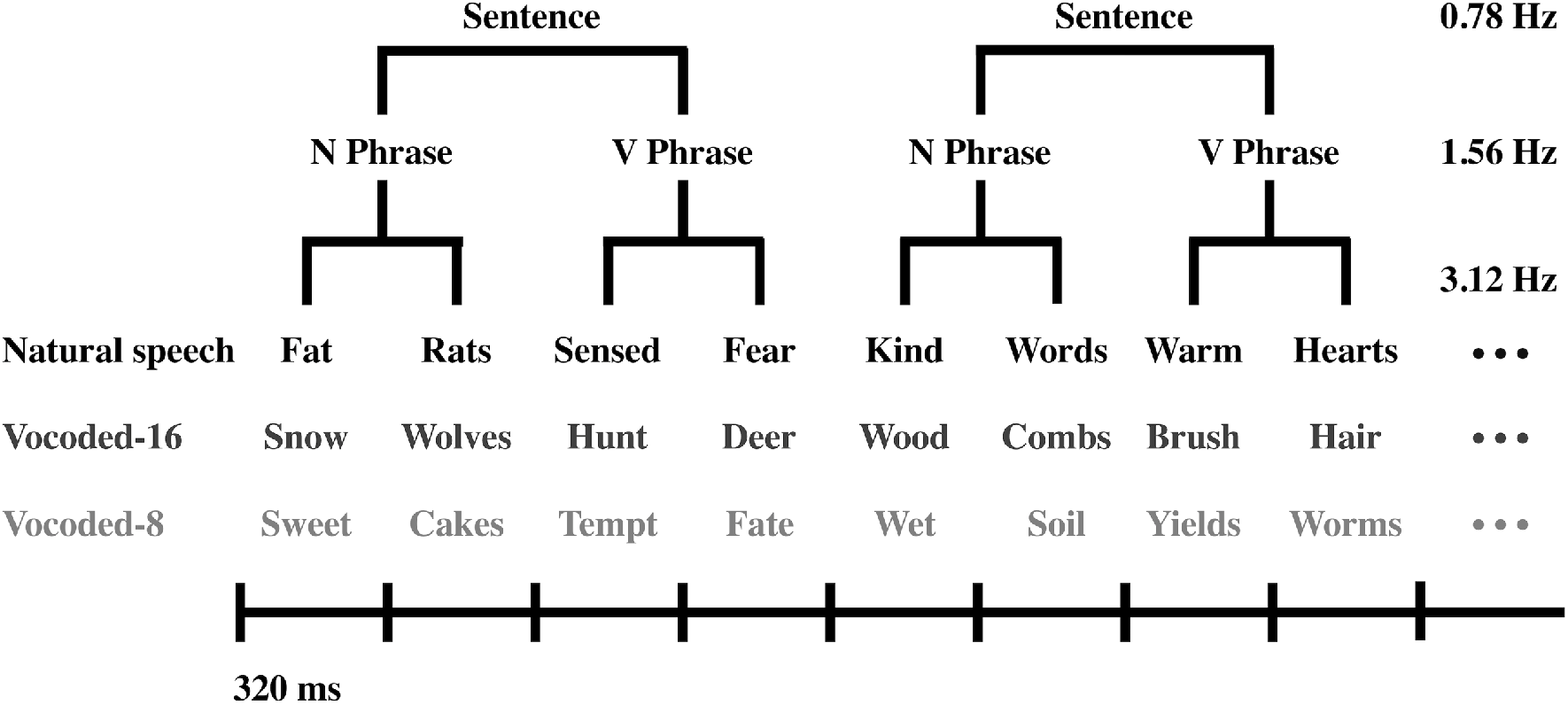
Sequences of English monosyllabic words under different intelligibility conditions were presented isochronously, forming phrases and sentences. N and V depict noun and verb, respectively.

### Noise Vocoding

The remaining 120 four-syllable sentences (after the selection of 60 sentences of “natural speech”) from the sentence list were processed with noise vocoding to degrade intelligibility. Noise vocoding was performed using custom Matlab scripts (MATLAB and Statistics Toolbox Release 2016b, The MathWorks, Inc., Natick, Massachusetts, United States). The frequency range of 200 Hz to 22,050 Hz was divided into 16 or 8 logarithmically spaced channels using a 6th order Butterworth filter. Sixty randomly selected sentences were used to produce the 16-channel vocoded speech and the remaining sentences were used for the 8-channel noise vocoding. More spectral detail is achieved as a function of greater number of frequency channels used in the vocoding. In each frequency channel, the envelope of the speech stimulus was extracted with full wave rectification and low-pass filtered below 300 Hz (2nd order Butterworth filter). This envelope was then used to amplitude modulate white noise filtered into the same frequency channel from which the envelope was extracted. These envelope-modulated noises were then recombined over frequency channels to yield the noise-vocoded speech segments. The root-mean-square (RMS) level of the noise-vocoded stimulus was normalized to match that of the original speech signal.

To validate and quantify the effect of the intelligibility manipulation, a behavioural word report task was performed by a separate group of native English speakers (n = 23; 18 −32 years old, mean 22 years old; 19 female). In this task, participants heard one four-syllable English sentence at a time and were required to type it out on the screen using a keyboard. As the shuffled speech sentences were designed to be entirely unintelligible, only the natural and noise-vocoded sentences were tested. Several example sentences were played to each participant first and then sentences from each condition were presented via headphones (Sennheiser HD 280 Pro) in separate blocks (order randomized across participants) at a comfortable listening level. Participants indicated they had finished typing a trial by pressing the return key, which initiated presentation of the next sentence with a delay at 1.2 s. Each block had 60 sentences and the sentences were presented in a random order. The percentage of correctly reported words is shown in **Figure 2**. As expected, accuracy for the natural speech condition was significantly higher than for the other conditions (voc 16: p < 0.001, voc 8: p < 0.001, paired one-sided t test) while the accuracy for the 16 channels noise vocoded condition was significantly higher than for the 8 channel condition (p < 0.001, paired one-sided t test).

**Figure 2:**
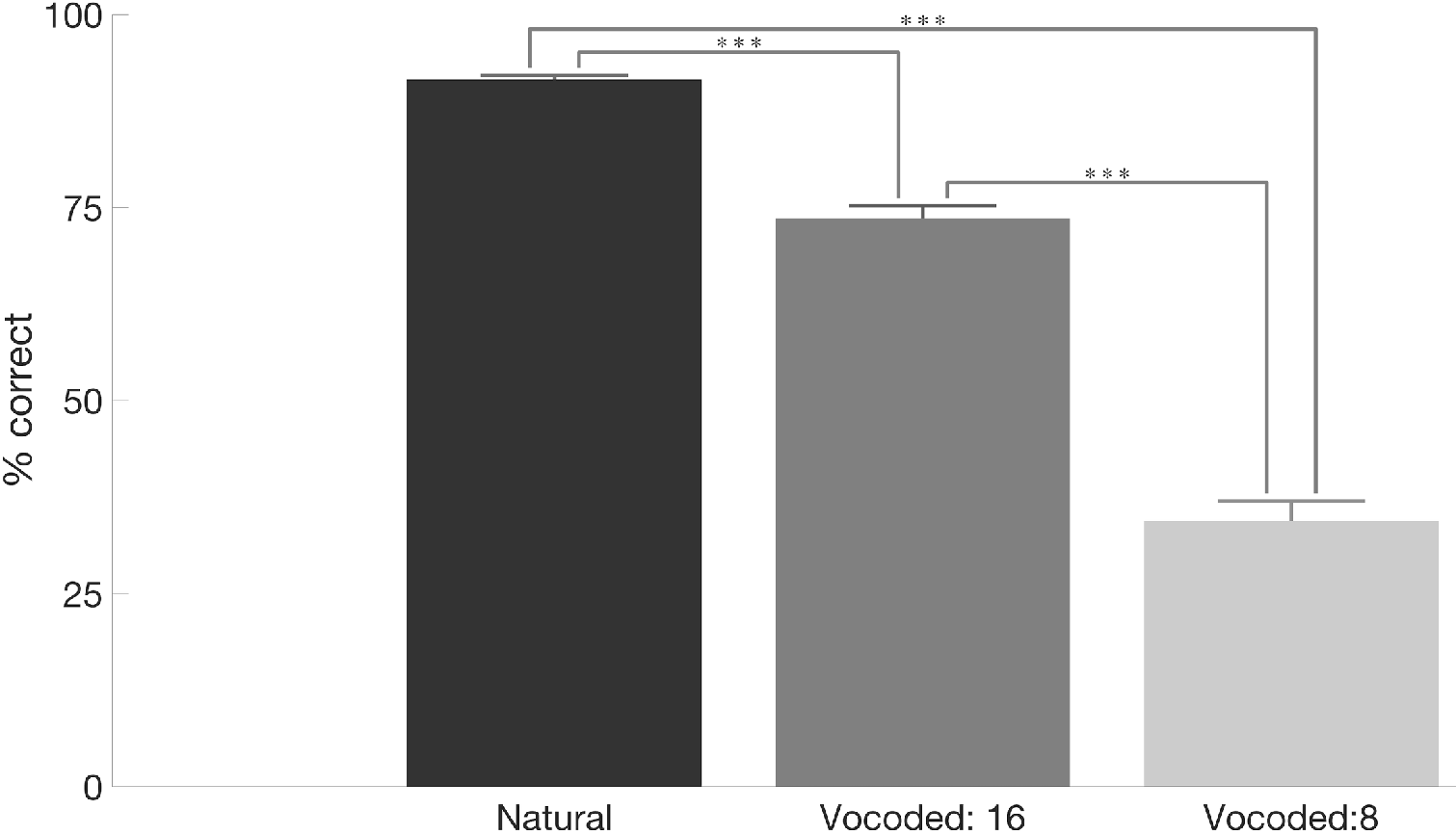
Performance of the word report task under different intelligibility conditions (natural, vocoded speech: 16 and vocoded speech: 8, error bars reflect ±SEM).

### Shuffled Speech

The same set of 60 four-syllable sentences used for the “natural speech” condition was employed again to produce shuffled sound streams as described in Ding et al. (2016). Each syllable in the original sentence was segmented into five overlapping slices of 72 ms in length and with 10 ms portions overlapping with neighbouring slices. The overlapped region for each slice was smoothed by a tapered cosine window, except for the first slice (onset) and the last slice (offset) of a sentence. As indicated by its name, a shuffled speech sentence was constructed by shuffling all slices at the same position across sentences so that the slices in a given sentence were all replaced by slices randomly chosen from different sentences at the corresponding position. In a shuffled speech trial, 12 different shuffled sentences were played sequentially and were the same length of the natural or noise-vocoded speech trials. In an outlier trial, four consecutively shuffled syllables were replaced by four randomly chosen monosyllabic English words from the sentence list of 60 that did not form a sentence (e.g. trim, fruit, tails, soap).

### Stimulus Characterization

Acoustic properties of the speech stimuli used were characterized by the slow varying temporal envelope which reflects the sound intensity fluctuations. The amplitude envelope for each stimulus was extracted using half-wave rectification and the mean power spectrum shown in **Figure 3**, was acquired by applying a Fast Fourier Transform (FFT) to individual amplitude envelope and then averaging within each condition.

**Figure 3:**
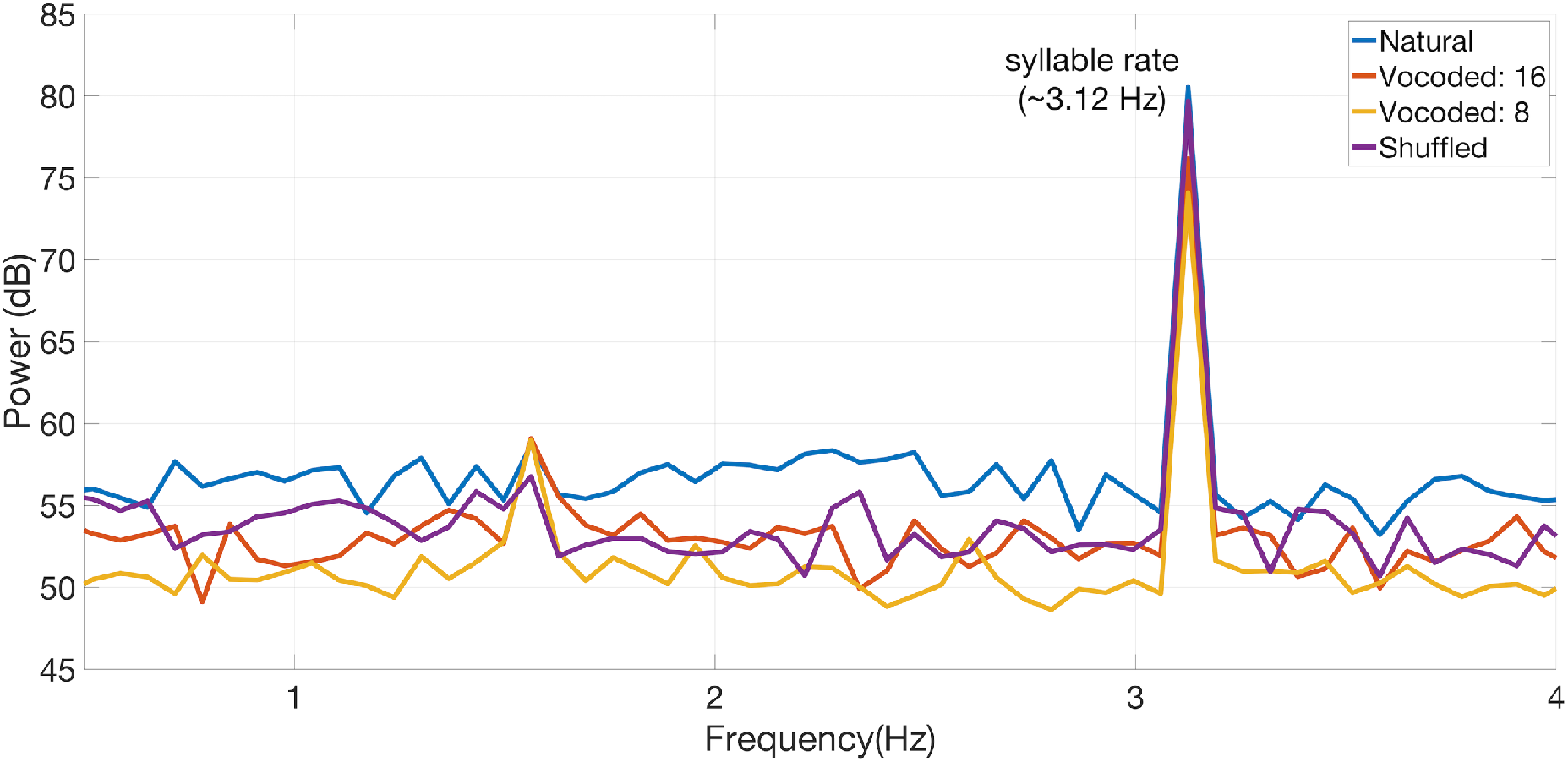
Power spectra of the different speech stimuli. Across the four intelligibility conditions, the stimulus power was all strongly modulated at syllable rate (~3.12 Hz) but not at phrase (~1.56 Hz) or sentence rates (~0.78 Hz) (see Figure 1).

### Experimental Procedure

Example sentences from each condition were played to each participant prior to the experiment. Natural four-syllable English sentences, noise vocoded four-syllable English sentences using 16 and 8 channels and shuffled sequences were presented in separate blocks at 75dB sound pressure level (SPL) through custom built, high fidelity insert earphones (Raicevich, Burwood, Dillon, Johnson, & Crain, 2010) with a flat frequency response up to 8 kHz. The order of the blocks was counterbalanced across participants. Participants were instructed to fix their gaze on a central cross projected to a ceiling screen and indicate whether it was a normal trial or an outlier trial via button press (index and middle finger, right hand) at the end of each trial. The button press initiated presentation of the next trial with a randomly-selected delay of 1.2 s, 1.4s or 1.6 s. Each block had 22 normal trials and 8 outlier trials and the trial types were presented in a random order.

### MEG & MRI Data Acquisition

Prior to MEG recordings, marker coil positions and head shapes were measured with a pen digitizer (Polhemus Fastrack, Colchester, VT). Brain activities to speech streams under different intelligibility conditions were recorded continuously using the KIT-Macquarie MEG160 (Model PQ1160R-N2, KIT, Kanazawa, Japan), a whole-head MEG system consisting of 160 first-order axial gradiometers with a 50-mm baseline (Kado et al., 1999; Uehara et al., 2003). MEG data was acquired with the analog filter settings as 0.03 Hz high-pass, 200 Hz low-pass, power line noise pass through and A/D convertor settings as 1000 Hz sampling rate and 16-bit quantization precision. The measurements were carried out with participants in a supine position in a magnetically shielded room (Fujihara Co. Ltd., Tokyo, Japan). Marker coils positions were also measured before and after each recording block to quantify participants’ head movement, the displacements were all below 5mm. The total duration of the experiment was about 45 minutes.

Magnetic resonance images (MRI) of the head were acquired for 19 participants at the Macquarie University Hospital, Sydney, using a 3 Tesla Siemens Magnetom Verio scanner with a 12-channel head coil. Images were acquired using an MP-RAGE sequence (208 axial slices, TR = 2000 ms, TE = 3.94 s, FOV = 240 mm, voxel size= 0.9 mm^3^, TI = 900, flip angle = 9°).

## Data Analysis

MEG data analysis was performed on normal trials only (excluding the catch trials), using the FieldTrip toolbox (Oostenveld, Fries, Maris, & Schoffelen, 2011) and custom MATLAB (The MathWorks, Inc., Natick, Massachusetts, United States) scripts. Offline MEG data were first filtered with a high-pass filter (0.1 Hz), a low-pass filter (30 Hz) and a notch filter (50 Hz, 100 Hz, 150 Hz) and then segmented into epochs according to trial definition. To avoid excessive stimulus-onset evoked responses, only the data between start of the second sentence (or the fifth syllable if the stimulus contained no sentential structure) and the end of each trial were analysed further. All data trials were down-sampled to 200 Hz prior to independent component analysis (ICA)(Makeig, Bell, Jung, & Sejnowski, 1996) to remove eye-blinks, eyemovements, heartbeat-related artefacts and magnetic jumps. Components corresponding to those artefacts were identified as by their spectral, topographical and time course characteristics. After ICA artefact rejection, all 22 cleaned trials of MEG data were averaged in the time domain.

### Sensor Level Analysis

Data analysis was carried out in the frequency domain to reveal brain activities tracking the different levels of linguistic units. Frequency spectra were calculated by applying FFT to the time-domain averaged MEG data (14.08 s) with a Hanning window, resulting in a frequency resolution of approximately 0.071 Hz.

A recent study by Zhang and Ding (2017) demonstrated that the tracking of hierarchical linguistic structures emerges at the beginning of the stimulus and are reflected by slow neural fluctuations, rather than a series of transient responses at boundaries (Zhang & Ding, 2017). Motivated by these time-domain characteristics, we also calculated the magnitude-squared coherences between the MEG recordings and a composite signal, which is comprised of sinewaves at frequencies corresponding to the presentation rate of syllables/words, phrases and sentences respectively (**Figure 4**). Magnitude-squared coherence is a frequency-domain measure of phase consistency between two signals across multiple measurements, with a normalized value between 0 and 1. MEG data trials, as well as the composite signal, were segmented into short frames of 2.56-second in length and transformed to the frequency domain with FFT and using a sliding Hanning window (50% overlap, 10 frames/trial, ~ 0.39 Hz frequency resolution). Coherence was then calculated with the power spectral density of each MEG channel and the cross-spectral density between each MEG channel and the composite signal, estimated from the frequency transformed data frames.

**Figure 4:**
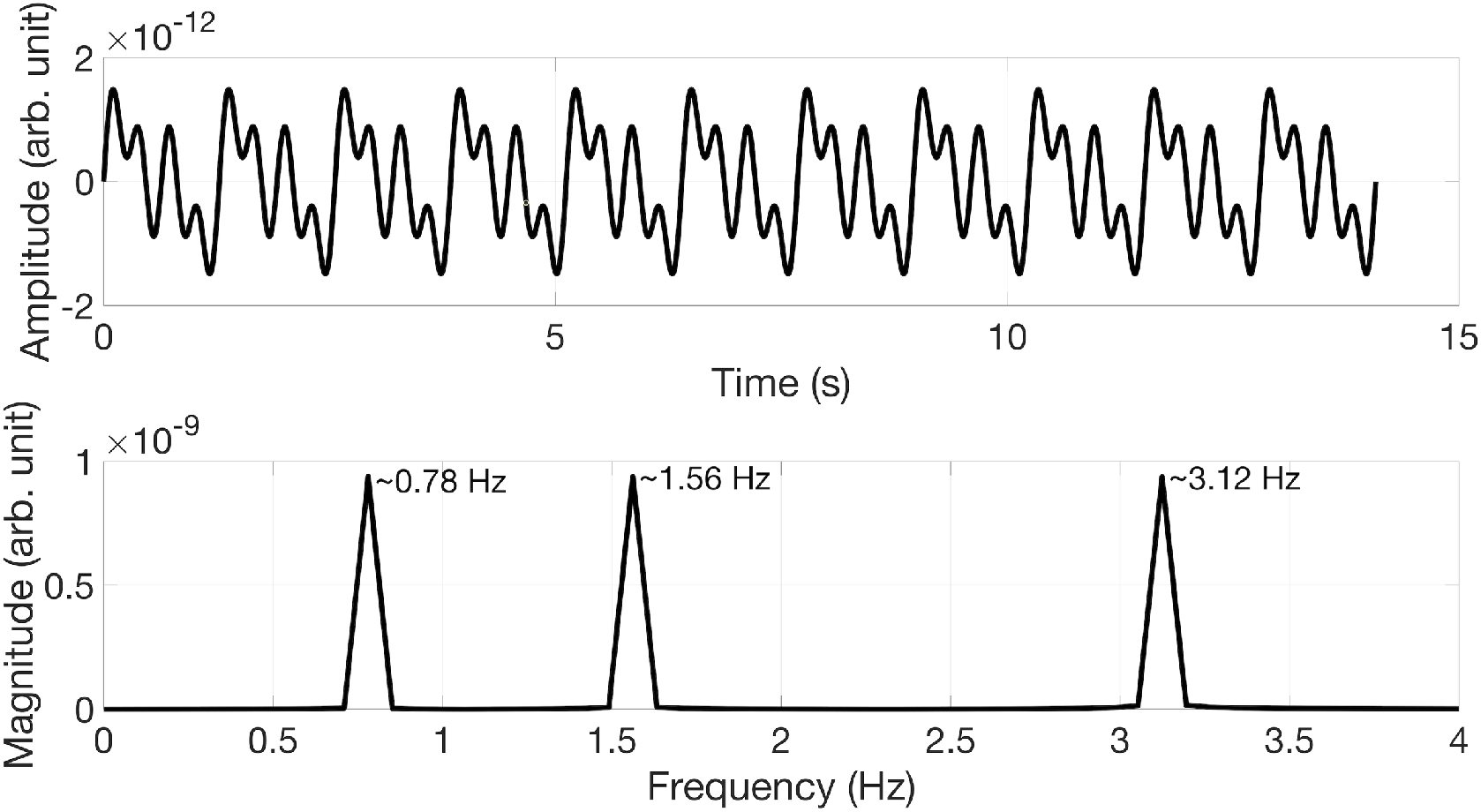
The composite signal used for coherence calculation. Top: time series of the composite signal with a duration equivalent to 11 short sentences. Bottom: Frequency domain representation of the composite signal shows peaks at sentence rate (~0.78 Hz), phrase rate (~1.56 Hz) and syllable rate (~3.12 Hz) respectively.

### Source Analysis

To investigate the spatial distribution of cortical areas coherent to different levels of linguistic structure, we conducted a whole-brain beamforming analysis using Dynamic Imaging of Coherent Sources (DICS) (Gross et al., 2001) which is a frequency domain, linearly constrained minimum variance beamformer (Veen, Drongelen, Yuchtman, & Suzuki, 1997). Source models were constructed based on each individual’s structural MRI. Cortical surface reconstruction (white-grey matter boundary) and volumetric segmentation was performed with the Freesurfer image analysis suite (Fischl, 2012); http://surfer.nmr.mgh.harvard.edu/). Cortical mesh decimation (ld factor 10 resulting in 1002 vertices per hemisphere) and surface-based alignment was performed with SUMA - AFNI Surface Mapper (Saad & Reynolds, 2012). A single shell volume conduction model (Nolte, 2003) was adopted and the 2004 cortical surface vertices were used as MEG sources for the leadfield calculation. For more details of the source head modelling procedure see Li Hegner et al. (2018).

DICS was applied to the FFT transformed MEG data frames at the corresponding frequency of each linguistic unit across all intelligibility conditions. Coefficients characterizing the beamformer were computed from the cross-spectral density matrix and leadfield matrix at the dominant orientation. Source level coherence images were generated by calculating coherence values between neural activity at each vertex (source point) and the composite signal using the resulting beamformer coefficients. Random coherence images were generated as the average of 100 source space coherence values calculated using the same composite signal but were randomly shuffled at each time, similar to the implementation described by Peelle et al. (2013). Cortical level group analyses were performed using cluster-based permutation tests to correct for multiple comparisons (Maris & Oostenveld, 2007) with a critical value of alpha = 0.01 and 1000 random permutations. Each coherence image was contrasted with a corresponding random coherence image; the effect of speech intelligibility was evaluated by contrasting coherence images across levels of intelligibility.

## Results

### Behavioural results

Error rates for the behavioural task (designed to ensure maintained vigilance) of the different experimental conditions during MEG data acquisition are summarised in Table 1. Error rates were calculated by averaging the miss rate and the false alarm rate under each intelligibility condition. Errors were significantly lower in the natural speech condition (voc 16: p = 0.016, voc 8: p = 0.001) whereas the accuracy for the two vocoded conditions did not differ from each other (p = 0.96), assessed by paired two-sided t tests. As the experimental task for shuffled speech condition was different (detect consecutive monosyllabic English words embedded in shuffled syllable streams), the behavioural performance was not statistically compared with the other conditions.

**Table 1:** Behavioural performance for all experimental conditions (error rate mean ± SEM)

| Natural Speech | Vocoded Speech: 16 | Vocoded Speech: 8 | Shuffled Speech |
| --- | --- | --- | --- |
| $36.8 \pm 2.3\%$ | $45.6 \pm 2.8\%$ | $45.7 \pm 1.5\%$ | $18.3 \pm 2.4\%$ |

### Phase-Locked Responses to Hierarchical Linguistic Structures

Frequency response and coherence with the composite signal calculated under different intelligibility conditions were grand averaged across all participants as well as all MEG channels and plotted in **Figure 5**. Compared with the averaged power spectra of speech stimuli (**Figure 3**), it is evident that both frequency response and coherence plots show peaks corresponding to the phrase rate (~1.56 Hz), sentence rate (~0.78 Hz) and syllable rate (~3.12 Hz). At the phrase and sentence levels, mean response magnitude and coherence both declined as a function of decreasing speech intelligibility. In contrast, the mean syllable level response magnitudes and coherence showed no systematic relationship to intelligibility levels.

**Figure 5:**
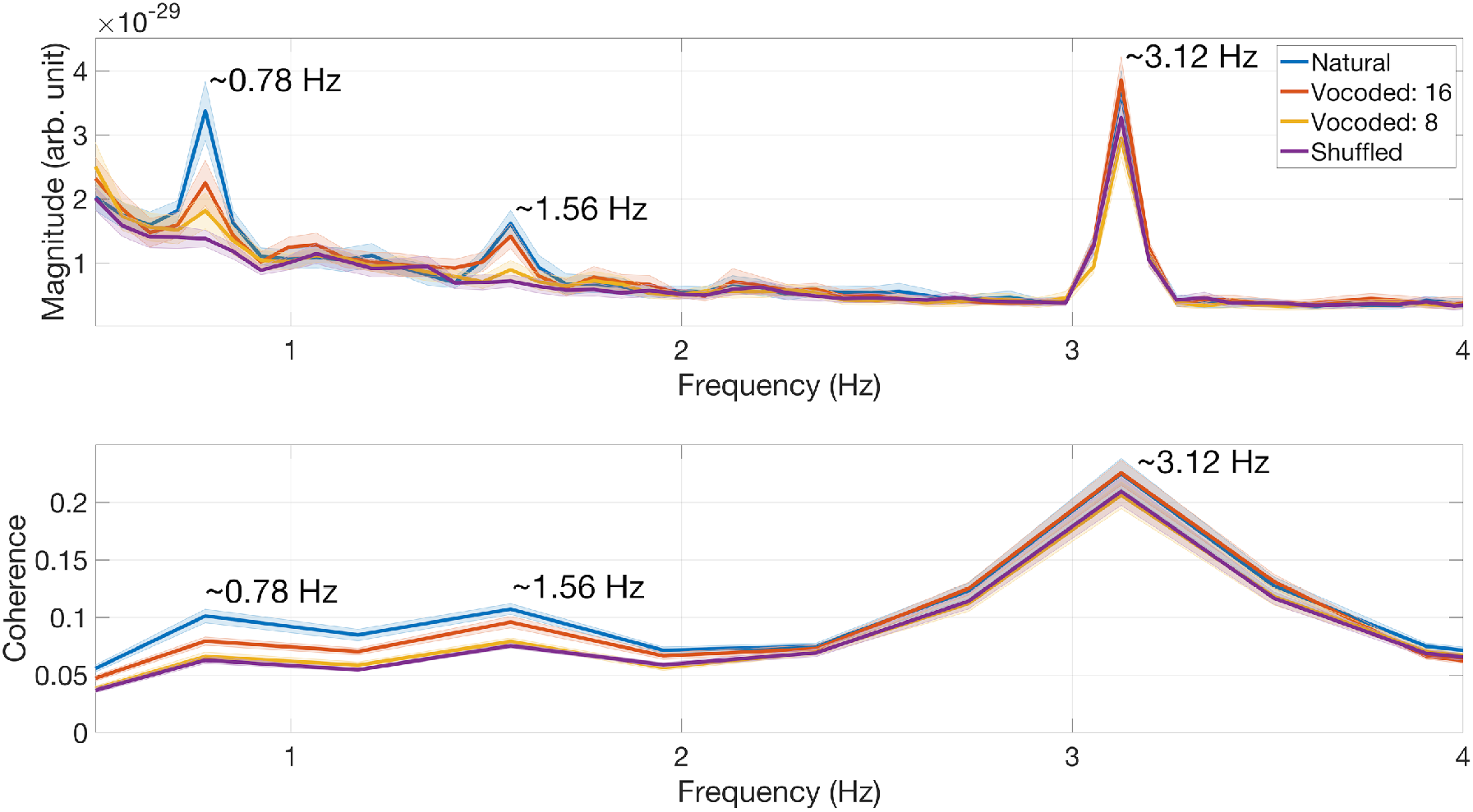
MEG tracking differs under different intelligibility conditions. Top: Averaged MEG sensor level frequency responses (160 channels)) exhibited different tracking activity to the hierarchical linguistic information (syllable, phrase and sentence). Bottom: Averaged MEG sensor level coherence between each MEG channel and the composite signal. The shaded area indicates 2 S.E.M.

### Cortical Sources Coherent to Hierarchical Linguistic Structures

The DICS source localization results (quantified as coherence values) were overlaid on the cortical mesh of each individual participant. For visualization purposes, source space results were grand averaged and plotted on a common brain mesh generated using the Freesurfer template brain (http://surfer.nmr.mgh.harvard.edu/), segmented and processed following the procedure described in the Data Analysis section.

**Figure 6** shows grand mean source coherence results for each experimental condition and linguistic unit. Several features are worth noting prior to statistical analyses. First, mean coherence at the syllable level was bilateral and similar in size, in both hemispheres, across all experimental conditions. Second, mean coherence values at the phrase and sentence levels were larger in the left hemisphere, and declined (in both hemispheres) as a function of decreasing intelligibility.

**Figure 6:**
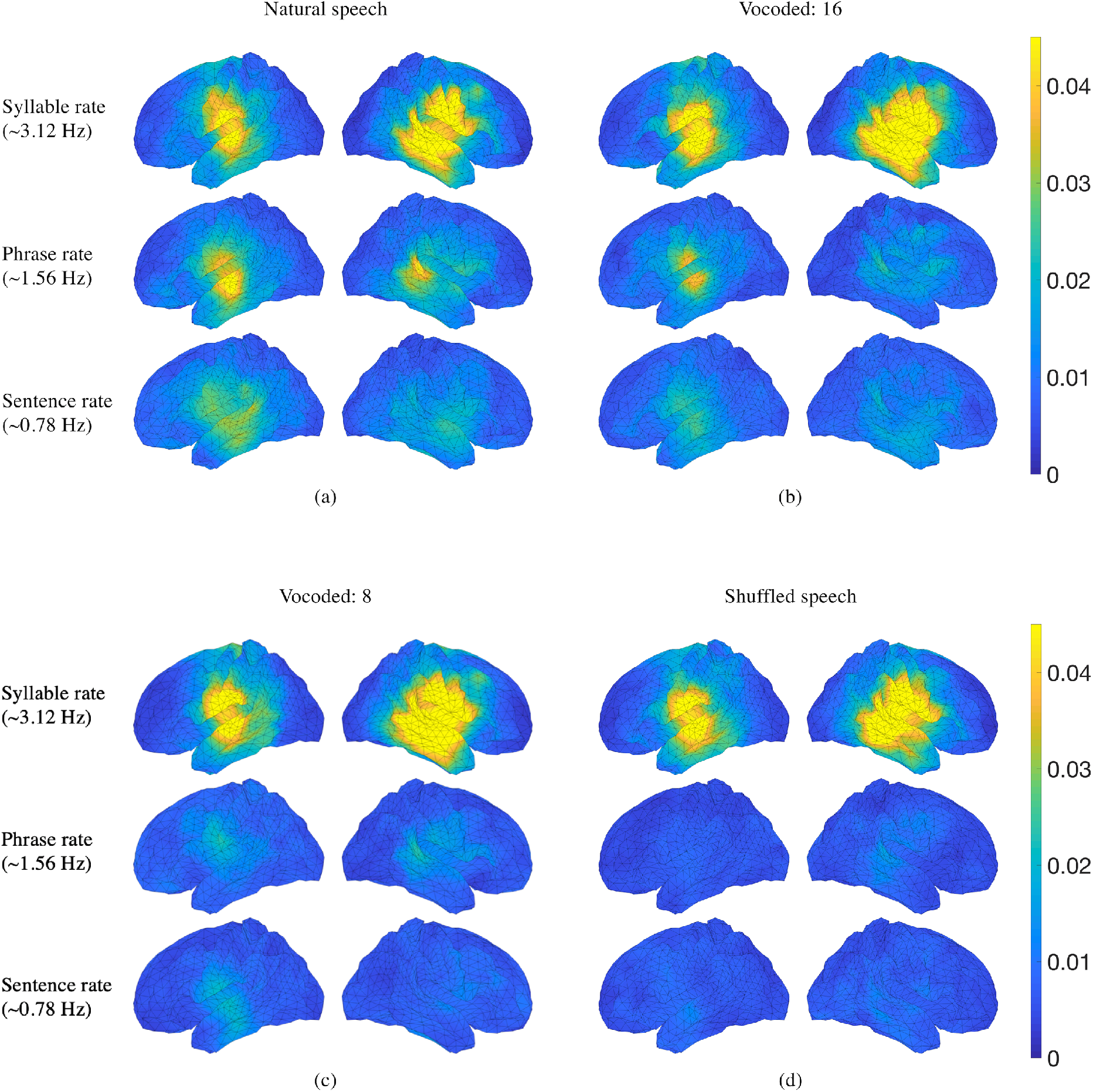
Source localization results grand averaged across all 19 participants and plotted on a template cortical mesh. Grand averaged coherence values at frequencies corresponding to syllable linguistic structure are sustained across all intelligibility conditions. (a): Grand averaged coherence values at frequencies corresponding to all three linguistic units are left-lateralized under natural speech condition. (b): Grand averaged coherence values at frequencies corresponding to phrase and sentence rates are reduced under 16 channel vocoded speech condition. (c): Grand averaged coherence values at frequencies corresponding to phrase and sentence rates are further reduced under 8 channel vocoded speech condition. (d): Grand averaged coherence values at frequencies corresponding to phrase and sentence rates are diminished under shuffled speech condition. Colour bars indicate coherence values.

### Contrasts with random coherence

Whole-brain analyses contrasted coherence maps in each experimental condition against “random” coherence maps (calculated using shuffled composite signals – see Methods section). Results are shown in **Figure 7** using paired one-sided t tests with a sample-wise threshold of *p* < 0.005 and a threshold of *p* < 0.005 whole-brain cluster extent multiple comparison correction.

**Figure 7:**
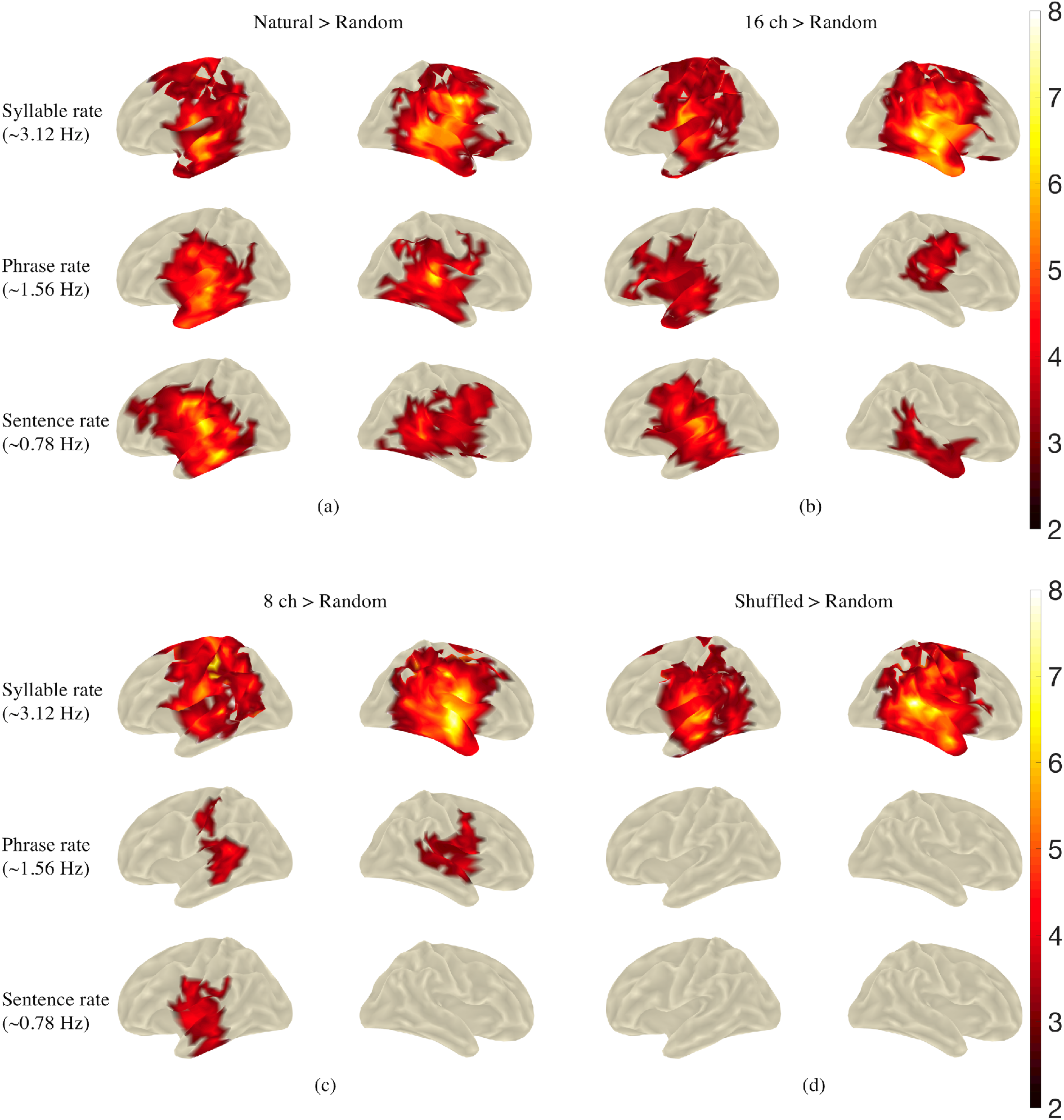
Contrasting source localized coherence tracking hierarchical linguistic structures under different intelligibility conditions with random coherence. (a) Cortical areas showing bilateral significant coherence at all three frequencies under natural speech condition. (b) Cortical areas showing bilateral significant but reduced coherence at frequencies corresponds to sentence and phrase rates under 16 channel vocoded speech condition. (c) Cortical areas showing further reduced (even unilateral) coherence at frequencies corresponds to sentence and phrase rates under 8 channel vocoded speech condition. (d) Cortical areas only showing significant coherence at frequency corresponds to syllable rate under shuffled speech condition. Colour bars indicate t values.

The results show that natural speech elicited significant coherence in regions surrounding bilateral auditory cortices, for all three linguistic structures. Notably, there were no significant vertices at the phrase and sentence rates for the shuffled > random condition, as would be expected since the shuffled condition was entirely unintelligible and therefore contains no information about phrase and sentence modulation. Another point to note is that the maps show greater coherence in the left hemisphere for the higher linguistic structures, especially at the lower intelligibility levels.

### Contrasts against shuffled speech

The foregoing contrasts provide a picture of the overall extent to which our measured neuronal responses tracked each of the three rates in the composite signal. In the next step, we wished to isolate neuronal responses to the abstracted linguistic units (phrase and sentence units) with contrasts against shuffled speech (which retains only the physical modulation at the syllable rate). In other words, the shuffled speech contrast allows us to remove the effect of any cues that are physically present in the speech stream.

As shown in **Figure 8**, positive significant clusters were found using paired one-sided t tests with a vertex-wise threshold of *p* < 0.005 and whole-brain cluster correction for multiple comparison at *p* < 0.005. No significant clusters were obtained for the syllable rate contrast (top row) or for the 8-channel contrast (not shown in the figure; presumably due to its low intelligibility). Notably, significant clusters for the sentence-level contrasts were restricted to the left hemisphere, as was the 16 channel phrase rate contrast.

**Figure 8:**
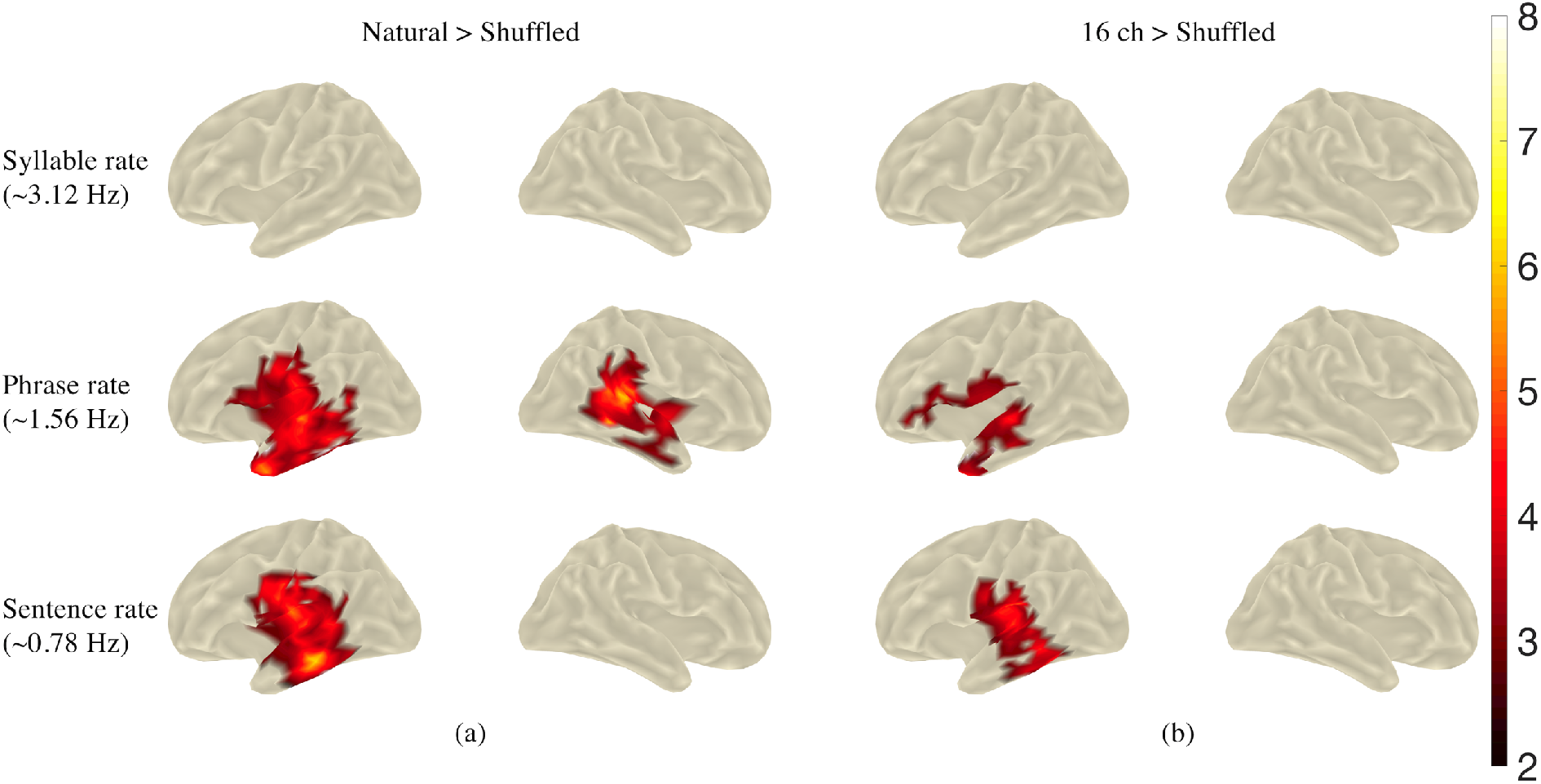
Contrasting source localized coherence tracking linguistic units across different intelligibility conditions. (a) Cortical regions showing enhanced coherence under natural speech condition compared to the shuffled speech condition. (b) Cortical regions showing enhanced coherence under 16 channel vocoded speech condition compared to the shuffled speech condition. No significant clusters were found for the 8 channel vocoded speech condition. Colour bar indicates t values.

## Discussion

The results of this study replicate those of Ding et al. (2016) and confirm that the human brain is sensitive to abstract linguistic structures in connected speech that are unambiguously dissociated from any acoustic cues to these structures.

The novel contribution of the present study is to demonstrate that the MEG responses to abstract linguistic structures are sensitive to parametric manipulations of speech intelligibility. The sensitivity was manifest in the data as: reduced coherence of MEG responses to embedded linguistic structures (phrase and sentence rates) as a function of reduced intelligibility; and coherent MEG responses to embedded linguistic structures became increasingly restricted to the left cerebral hemisphere as a function of decreased intelligibility. This lateralisation is particularly apparent at the sentence level.

The concurrent tracking of hierarchical linguistic structures demonstrated that brain speech processing is not only restricted to stimulus level analysis, i.e. a phenomenon which has been reported as cortical “entrainment” to speech temporal envelope at syllabic rhythm in many research works (Millman et al., 2015; Ding & Simon, 2014; Peelle, 2012; Luo & Poeppel, 2007). These multi-timescale brain dynamics provide a plausible neural mechanism of linguistic processing during speech perception through which smaller linguistic units can be temporally integrated into larger linguistic structures and eventually leads to comprehension (Ding et al., 2016). Our MEG data shows that brain responses synchronized to higher level linguistic structures are enhanced under intelligible speech condition when linguistic information is available, separated from speech acoustic signal processing. This suggests a potential neural framework for speech intelligibility during online speech processing and language comprehension. Furthermore, the same neural substrates may be shared during selective listening in a multi-talker environment, like the classic “cocktail party problem” (Cherry, 1953). Through the hierarchical linguistic structure building operation, a robust and distinctive auditory object is constructed from the target speech signal whereas unattended background speech streams are only minimally processed mainly to register syllabic rate fluctuations physically presented in the acoustic signal (Rimmele et al., 2015; Ding et al., 2014; Ding & Simon, 2012).

It is notable that intelligibility had little apparent effect on brain responses at the syllable level. The lack of effect on syllable level responses may be attributed to the fact that these are associated with responses evoked by the physical onsets of each syllable. The temporal envelope that reflects the energy fluctuation aligns with syllable rhythms in speech signals and were closely preserved across different intelligibility conditions with noise vocoding. This confound between acoustic and linguistic cues is inevitable in studies that employ naturalistic sentences for experimental stimuli (Zoefel & VanRullen, 2015). The results of the present experiment indicate that brain responses to mixed acoustic and linguistic cues may be largely driven by the acoustic cues and, as such, are relatively insensitive to manipulations of intelligibility. Such a lack of sensitivity may account for the mixed results reported in some recent studies of cortical “entrainment” to the envelope of speech signal under different intelligibility conditions (Ding et al., 2014; Ding & Simon, 2014; Howard & Poeppel, 2010; Millman et al., 2015; Peelle & Davis, 2012; Peelle et al., 2013; Zoefel & VanRullen, 2015). Importantly, and in contrast, our results indicate that unconfounded linguistic cues are clearly sensitive to intelligibility manipulations.

The lateralised results of our source analyses are striking. In his review of fMRI studies of speech comprehension, Peelle (2012) concluded that cortical lateralisation depends in a graded fashion on the level of acoustic and linguistic processing required. Processing related to non-speech signals (including amplitude modulated (AM) noise) is bilateral. As the requirements for linguistic analysis and integration increase (from AM sounds, through to phonemes, words, phrases and sentences), neural processing becomes increasingly left-lateralised. The present results are entirely consistent with this framework.

The noise vocoding employed in the present study mimics the sound processing strategies employed by cochlear implant devices (Shannon et al., 2004). Achieving intelligible speech is the major objective of this intervention. As such, the present results, demonstrating that MEG brain responses are clearly sensitive to intelligibility manipulations, taken together with the demonstration by Ding et al. (2017) that brain responses to abstract linguistic structures are also measurable with EEG recordings, indicate that these electrophysiological brain responses may serve as highly useful and objective neural markers of speech perception in cochlear implant recipients.

## Author Contributions

Q.M., Y.L.H., I.G., C.M and B.J. conceived and designed the experiment. Q.M. performed the MEG experiments. QM and Y.L.H performed data analysis. QM, Y.L.H and B.J wrote the paper. Y.L.H. and B.J. contributed equally as senior authors. All of the authors discussed the results and edited the manuscript.

## Funding

This work was supported by the Hearing Cooperative Research Centre (HearingCRC XR1.1.3) and Australian Research Council (grant number DP170102407). The authors also thank Dr. Nai Ding for his helpful discussion during the design of this experiment, Mr. Craig Richardson and Dr. Jessica Monaghan for their help with speech material generation.

## Notes

Conflict of Interest: None declared.

