## Supplementary Material for "Lateralised cerebral processing of abstract linguistic structure in clear and degraded speech"

### Supplementary Material: Sentence Stimuli

| | | |
|---|---|---|
| Fat rats sensed fear | Kind words warm hearts | Young kids close gates |
| Stacked shelves hold cans | Long fights cause hate | Flax threads hang plates |
| Big men drive trucks | Dead sharks spout blood | Their store sold jeeps |
| Bright flares shine light | Shrewd dogs dig holes | Wise cubs sip milk |
| Dry fur rubs skin | Lean girls like jeans | Four farms found cows |
| Sly fox stole eggs | Sick boys fail tests | Sharp knives cut cheese |
| Top chefs buy beef | Rear gates stop draughts | Soap suds cleanse toes |
| Our boss made deals | Firm palms make bread | Loud sounds scare moms |
| Two groups plant shrubs | Bad smells fill town | Weird clowns wear hats |
| All moms love kids | His aunt tied shoes | Her sons paint walls |
| New plans give hope | Quiet lambs graze grass | Giant bears walk trails |
| Large ants built nests | Soft forks spill rice | Drunk dudes sang tunes |
| Teen apes chase bugs | Tree frogs stalk flies | Small chicks catch grubs |
| Rude cats claw dogs | Black skies show stars | Brown bags take space |
| Rich cooks brew tea | Tall guys flee camp | Hot grills cook steaks |

| | | |
|---|---|---|
| Fun games waste hours | Grey sheep seek hills | Big rocks clog roads |
| Pink toys please girls | Iced beer costs bucks | Storm floods ruin farms |
| Great waves wreck ships | Brave kings fight wars | Warm ground melts snow |
| Vain ears hear talk | Sore eyes shed tears | Keen blades slash tires |
| Close friends swap gifts | Harsh trails sprain joints | Posh wives pay bills |
| Horse hooves crush rocks | Mad dogs bite tails | Good shops pour drinks |
| Red lights stall cars | Fine gifts please hosts | Some pets climb trees |
| House maids scrub floors | Fierce flames sear steak | Snow limbs lift weights |
| Oil lamps start fires | Snow wolves hunt deer | Chrome tanks leak gas |
| Wood combs brush hair | Sheer noise hurts ears | Smart girls read books |
| Cold storms harm plants | John’s wife bakes cakes | South lane leads home |
| Bowled balls strike pins | Blunt sticks smash glass | Parched fields need help |
| Three teams lost games | Smooth eggs hatch chicks | Steep hills slow bikes |
| Gas stoves heat pans | Cute birds build nests | Deep trust bonds friends |
| Cheap baits halt slugs | Sour food draws breath | Low planes dust crops |
| Weak sun heats rooms | Tight fists knock doors | Cruel hooks catch fish |

| | | |
|---|---|---|
| Bee stings prick arms | Rough walks tax legs | Thick fog blocks views |
| Straw hats stop sun | Chopped logs choke creeks | Fire doors seal smoke |
| Farm aids swing bats | Tough spades crack slabs | Bald men ride trains |
| Old pumps lack grease | Spare keys lock halls | Plump wool coats sheep |
| Stretched arms seize balls | Iron spoon knocked floor | Wall clocks tell time |
| Round box stores coins | Bank clerks scan files | Square nets grab prawns |
| Dried fruit tastes good | Deep breaths save life | Mild rain wets ground |
| Gold rings cause fights | Small hands knead dough | Slim hips twirl hoops |
| Shoe tread stops slips | Thin ice risks lives | Brick walls guard homes |
| Fried chips burn tongues | Dark nights veil owls | Dear friends send mail |
| Open sports draw crowds | North winds bring joy | Sweet cakes tempt fate |
| Tall trees lose leaves | Lost goats scale cliffs | Grown men miss youth |
| Cracked plates spoil food | Wild pines drop cones | Spiked pins pierce rags |
| Chilled sheets help sleep | Bored dads drink beer | Wet soil yields worms |
| Bleak seas hide crabs | Bar soap cleans paws | Flat screws fix lights |
| Back teeth hurt jaws | Cool rooms keep meat | Calm swells raise yachts |

| | | |
|---|---|---|
| Huge bull jumps fence | Bus tours tire guests | Tight belts hold pants |
| Race cars dodge oil | White sand covers boats | Road bikes skip holes |
| King crabs eat shrimp | Clay mugs store pens | Short talks blow minds |
| Wine grapes have seeds | Aged trains use coal | Hedge plants block paths |
| Quick gales break kites | Hard falls hurt knees | Spring buds prize soil |
| Rose tea stains pots | Used bricks fill yards | Toy spoons stir cups |
| Hot baths treat flu | Green plants feed birds | Bush snakes kill mice |
| Axe strokes trim rope | Strong light fades cloth | Wide trucks move trash |
| Nice guys give seats | Silk scarves ease throats | Eight ducks cross fields |
| Brass clips grip notes | Blue pens write words | Long waits bore boys |
| Salt lakes ooze slime | Trust funds hoard wealth | Stiff brooms sweep stones |
| Fresh staff like work | Flight crews serve lunch | Steel whisks whip cream |
| Sun glare burns eyes | Clear tape seals splits | High racks hold coats |
